# Hot spots in a cold river synchronize temporarily separated salmon runs in their offspring’s development

**DOI:** 10.1101/2023.11.07.565965

**Authors:** JN Negishi, N Morisaki, YY Song, N Aruga, H Urabe, F Nakamura

## Abstract

The identification of thermal heterogeneity in the environment and its inclusion in adaptive strategies are key to habitat management of cold-water fish, including salmonid species. This study tested the hypothesis that upwelling of groundwater (GW) from a tributary catchment through a relatively deep aquifer (tributary GW) affects salmon redds selection in an urbanized gravel-bed river, and that the spawning preference of such areas depends on the seasonal context. The field study was conducted between 2001 and 2015 in an approximately 6-km long segment of the Toyohira River, Northern Japan. Chum salmon redds distribution data over 15 years (2001-2015) were combined with spatial distribution data of hyporheic water affected by tributary GW in the riverbed to examine seasonally variable redds site selection in relation to the presence of unique GW. Furthermore, models that predicted the hyporheic water thermal regime were coupled with redds count data to estimate the approximate timing of fry emergence from the riverbed. The redds site selection was seasonally variable, with a higher dependence on tributary-GW-affected areas with a decrease in water temperature. The time until fry emergence from spawning was shortened when the tributary-GW area was chosen during the cold winter. Overall, the present study identified hotspots for salmon spawning redds in winter with a disproportionately high level of site selection because of their warmer temperature compared to surface river water in winter. Thermally diverse spawning habitats allow the diversification of spawner strains in synchronized descents to the sea. Signs of tributary-GW pollution was suggested, and thus the conservation of the groundwater pathway and its sources, followed by improvements in quality, can be beneficial to the Chum salmon populations in the Toyohira River.

## 1. Introduction

In light of global environmental changes and ongoing natural degradation on a global scale, the protection, restoration, and maintenance of mechanical processes that dictate the distribution and abundance of diverse organisms in an adaptive manner is crucial for effective nature conservation (Millar et al., 2007). To minimize the anticipated impacts on current biodiversity in this context, understanding species sensitivity is key, which is only obtainable based on an accurate assessment of habitat use and how they are limited by the projected changes in specific environments (Williams et al., 2008).

Temperature regime changes are considered extrinsic stressors because many ectotherms, including freshwater species, are characterized by specific thermal adaptations and tolerances (Deutsch et al., 2008; Sunday et al., 2014; Comte et al., 2014). To this end, the identification of thermal heterogeneity and spatially temporally projected population dynamics of concerned species has progressed in recent years, and potentially available adaptive strategies have been proposed in contemporary freshwater research, especially for cold-water fish, including salmonid species (Fullerton et al., 2017; 2018; Morelli et al., 2020; Ishiyama et al., 2023). These relationships are spatially diverse and specific to underlying climates, geology, land use, and the level of hydrological alterations. Thus, more studies are needed to formulate effective adaptation strategies.

Many Salmonidae fish species have anadromous life cycles of ascending to upstream of rivers (salmon runs) in the summer to autumn seasons to spawn (lay their eggs), and their population viability is a concern (e.g., Jonsson & Jonsson, 2009; Kaeriyama et al., 2014; Crozier et al., 2021). Salmonidae spawning habitat (redds) quality has been extensively studied in relation to various environmental variables, including water temperature, as this phase governs the initial survival rates of populations. The eggs are laid in riverbed sediment, hatch as alevins, and remain buried in sediment until fry emerge in surface water. The zones where the egg to alevin stages are exposed correspond to the hyporheic zone (HZ) where groundwater and surface mixes (Boulton et al., 1998). Several key physicochemical environments of the riverbed and HZ are known to play a role in reducing site quality and preferences at the local scale because of their decisive effects on redd formation capacity of fish and fish mortality. First, fish have specific preferences for substrate-size materials that are dependent on body size (Kondolf, 2000). Second, HZ water quality is known to affect egg survival, especially through dissolved oxygen (DO), in terms of survival rates (Malcolm et al., 2003), which is related to particle size composition and porosity (Wu, 2000). HZ water quality is different from that of surface water, and the differences dynamically fluctuate in relation to seasonal and hydrological contexts, such as hydraulic head differences between HZ and surface water (Malcolm et al., 2004; Soulsby et al., 2009). As sediment particle size composition and HZ flow at the local scale are geomorphically controlled, redd sites have been associated with locations relative to channel bedforms.

HZ temperature is not a strong direct determinant of survival rates at the early life-cycle stage of salmonids, but its importance lies in its effects on developmental rates, timing of hatching, and emergence (Geist et al., 2006; Whitney et al., 2014). The timing variation in relation to temperature conditions (thermal regime) has been reported in the field among watersheds (Lisi et al., 2013) and segments longitudinally located from upstream to downstream within a river >10 km from each other (Webb & McLay, 1996). HZ water temperature differs at various spatial scales under the control of surface water and groundwater influx (Weigelhofer & Waringer, 2003; Alexander & Caissie, 2003; Wu et al., 2020; Marmonier et al., 2020). The conservation of such diverse thermal heterogeneity is potentially important because the developmental speed and timing of emergence and descent to the sea reflect an adaptation to preferable downstream conditions, including the sea. For example, eggs laid in warmer HZ can theoretically offset the costs associated with delayed spawning timing compared to eggs laid in colder HZ in earlier seasons by shortening the development period. Furthermore, recent studies have demonstrated that such thermal heterogeneity manifested by thermally distinct habitats via groundwater or shallow upwelling flow can mediate site selection of redds, even at the scale of approximately 10 km (Mouw et al., 2014; Aruga et al., 2023). These studies highlighted the importance of superimposing thermal conditions onto other known physical-chemical variables to accurately account for the mechanistic understanding of redds formations. However, no study has quantitatively estimated how such thermal HZ habitat diversification can be reflected in egg development rates and fry descent timing.

This study tested the hypothesis that upwelling of groundwater (GW) from a tributary catchment through a relatively deep aquifer (tributary-derived GW) affects salmon redds selection in an urbanized gravel-bed river, and the spawning preference of such areas depends on seasonal context. Groundwater pathways in gravel-bed river systems are spatially multiple, ranging from relatively deep valley-scale flow to relatively shallower and shorter flows, as well as from tributary catchments (Power et al., 1999; Geist & Dauble, 1998; Marmonier et al., 2020). In this study, we focused on the relatively deep groundwater from the tributary catchment (Negishi et al., under review) because no previous studies have focused on this GW type in fish spawning site selection. We predicted that GW forms a HZ with warmer water temperature relative to surface river water in winter when the surface water temperature decreases, and that redds selection becomes disproportionately skewed to the GW-affected HZ in winter. This study is an extension of a previous study by Aruga et al. (2023) in the same river in which the selection of redds in relation to geomorphic units differed among seasons with the suggested effects of GW. We focused on Chum salmon, Oncorhynchus keta, a numerically dominant anadromous salmon in the study area, Toyohira River, Hokkaido, Japan. We first identified a section of GW-affected HZ based on multiple physicochemical parameters of water, followed by the analyses of redds distribution in relation to HZ characteristics in different winter months. To determine the importance of thermally diverse HZ water associated with GW in mediating salmon population dynamics, the HZ water temperature was modelled to simulate emergence timing differences with and without the GW-affected HZ. A detailed understanding of the processes and mechanisms leading to the seasonally dynamic spatial distribution of Salmonidae redds and its potential consequences can provide guidelines for GW regulations aimed at fulfilling the biological and hydrological requirements for riverine ecosystems in changing environments.

## 2. Material and methods

### 2.1 Study site

The field study was conducted between 2001 and 2015 in an approximately 6-km long segment of the Toyohira River (Fig. 1), which flows through Sapporo city and is a tributary of the Ishikari River with 72.5 km long and has a drainage area of 902 km^2^. This segment currently provides >95% of the Chum salmon redds sites in the Toyohira River and encompasses the area affected by groundwater (Aruga et al., 2023). The segment located in the lower part of the Toyohira Alluvial Fan and river water is affected by several multi-purpose dams in upstream areas, and surface runoff and groundwater affected by the urbanized area whose population was ca. 380,000. Although the natural river landscape of the studied segment was typical of gravel-bed rivers with well-developed gravel bars and multiple braided channels, installations of embankments and groundsills and channel straightening over the past 100 years transformed the river into one limited lateral channel migration. Despite such substantial environmental changes, several hundred to a few thousand Chum salmon redds are formed annually (Aruga et al., 2023). In the study segment, groundwater originating from a tributary catchment, which has been lost over landscape transformation in the last 60 years, continues to form characteristic hyporheic water (high in nitrate and low in dissolved oxygen) (Negishi et al., under review). Benthic organisms and riparian invertebrates were more abundant in sections affected by this characteristic groundwater. The mean annual flow rate and mean annual maximum flow rate over the past several decades were 26 and 440 m3/s, respectively. The river has a typical snowmelt hydrograph – high flow in late spring and early summer and low flow in late summer and winter, with water temperature being highest in July to September at >15 °C and lowest in January and February at <3 °C (Supplementary material 1).

**Figure 1.**
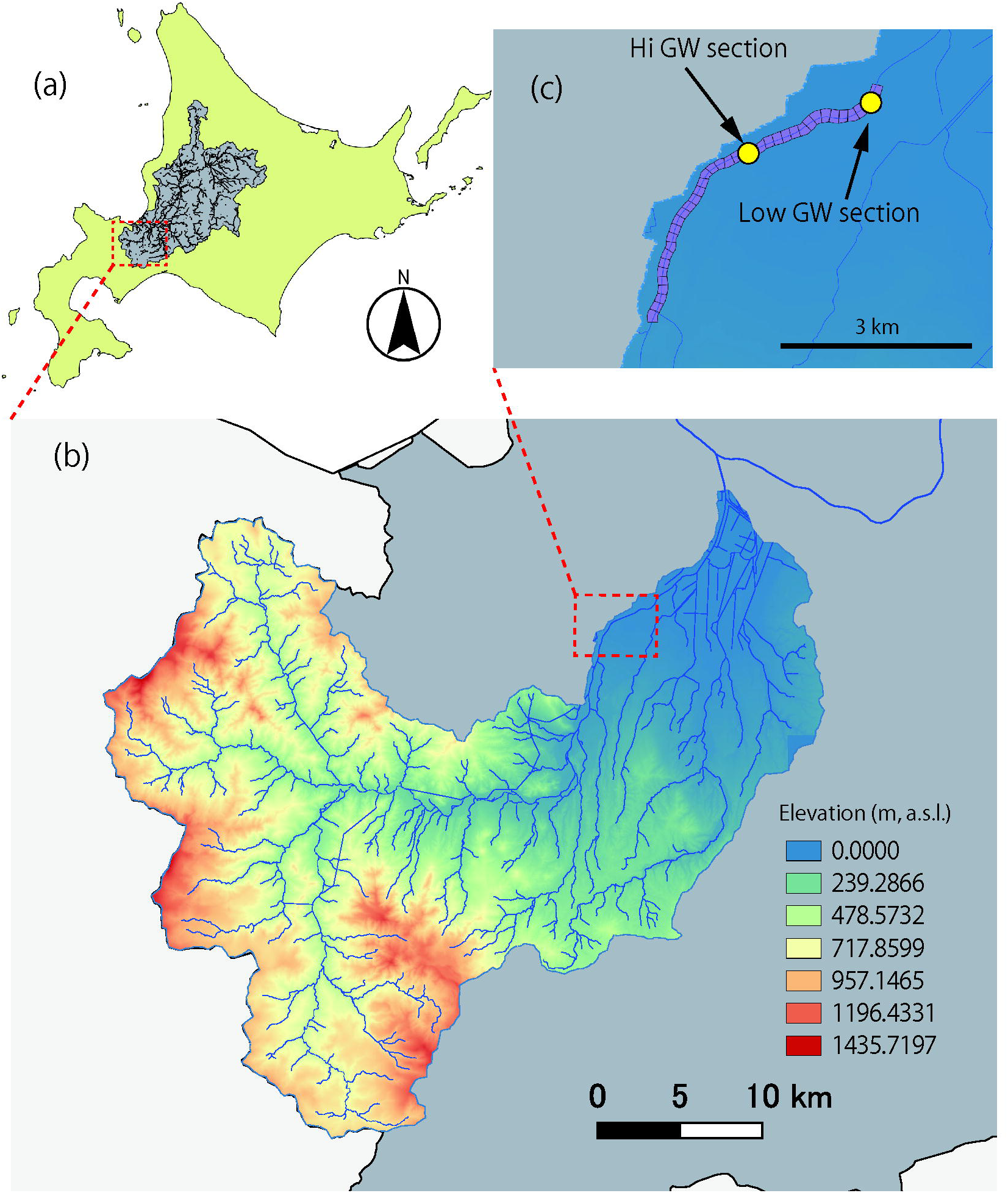
Map showing the study area in the Ishikari River watershed system (a), Toyohira River watershed (b), and the study segment (c). In (b), color gradation indicates elevation (m above sea level), with a channel network in blue lines. In (c), the purple-colored polygon shows 150-m compartments of the study segment used to organize data (see the text); yellow circles denote 250-m sections used for intensive monitoring surveys.

### 2.2 HZ water characterizations

Measurements were conducted in two ways: seasonal snapshot longitudinal measurements across the section and intensive continuous measurements in selected sections (Fig.1).

Snapshot measurements were conducted both in winter and summer (29-30 March and 17 August, 2012). The study segment was walked down along the channel thalweg, and measurements were taken at spots every 50 m (Negishi et al., under review). To represent diverse potential redds sites in river channels, at each distance, spots were chosen regardless of habitat type, and the water depth was <60 cm in pools but never <20 cm; 60 cm was the maximum depth that our methods can take measurements. At each chosen spot, a 1-m long stainless-steel rod (inner diameter 9 mm) was pounded into the riverbed to a depth of 25 cm, a perforated (lower 5 cm) acrylic tube was inserted into the rod, and the rod was removed so that the perforated tubes were buried at a depth of approximately 15 cm from the riverbed surface.

Thermocouples connected to a thermometer (TX 10, Yokogawa Meter & Instruments, Co.) were inserted into the tube to measure the water temperature, followed by pumping up approximately 500 ml of HZ water to measure the dissolved oxygen (DO), electrical conductivity (EC), and pH using probes (WM-32EP, DKK-KOA Co. & HQ 40 d, HACH Co.). After measuring the HZ water, the surface river water was also measured for the same set of parameters. In August, owing to a rain event, a downstream section of 450 m was not measured.

Intensive monitoring was conducted in both winter and summer: February 6 for three weeks and August 9 for four weeks in 2016. Two 250-m sections, one each with and without tributary-derived GW inputs, were selected based on the abovementioned snapshot surveys (Negishi et al., under review). In each, five sub-sections with three sub-sites (10, 25, and 40 m from the upstream ends) were set up. At each sub-site, HZ water was collected and measured for the same set of parameters on the first day of monitoring, as was done for snap-shot surveys. Simultaneously, within 50 cm of the HZ water sampling point, temperature loggers (Hobo Water Temp Pro UA-001, Onset Co., Massachusetts, USA) were set at a depth of 15 cm from the riverbed surface to measure the HZ temperature every hour. This setup was conducted by pounding a steel pipe (internal diameter of 45 mm) with detachable plastic pointed heads to the desired depth, followed by setting the logger at the inside bottom end of the pipe and removing only the pipe so that loggers were left in the sediment with pointed heads. The loggers recovered after three– four weeks. Two additional loggers were set up in the surface water in each section.

### 2.3 Salmon redds survey

The fish survey was conducted as a routine environmental monitoring activity of the Sapporo Salmon Museum at least once per month and was increased to 2-3 times per month when possible (see Aruga et al., 2023). On each occasion, the individuals walked down along the channel and recorded freshly spawned redds of Chum salmon. Within the interval time between consecutive surveys, fresh redds became indistinguishable in terms of coloration because of the growth of periphyton; thus, these counts were considered independent and gave conservative estimates of redds abundance. Count data from to 2001-2015. For–2001-2010, the approximate locations of redds were recorded graphically on the site map, whereas the locations of redds were recorded using GPS (Map62SJ, Garmin Co.) in recent years. Data from to 2001-2010 were digitized and converted into GPS coordinates. Because of the uncertainty in the exact locations of digitized data (potentially spatial errors of >50 m) and compromise against data abundance, we compiled all data by counting in each 150-m long survey compartments at each survey occasion using QGIS version 3.16.6 (QGIS Development Team, 2021).

### 2.4 Other environmental factors

Two physical conditions were considered potential limiting factors in the selection of the longitudinal redds site. Cross-channel structures, such as check dams, are known to affect distribution and migration (Marschall et al., 2011). Stream gradients are also known to affect salmonid distribution (Burnett et al., 2007). River channel gradient (hereafter, channel gradient) was obtained for each 150-m compartment on regular channel cross-sectional profile elevational data of river channels by the Ministry of Land Infrastructure, Transport, and Tourism (MLIT) in 2011. There were five groundsills with fish ladders within the study segment for channel stabilization. The distance of each compartment from the immediate upstream groundsills (hereafter, GS distance) was obtained. Additionally, daily average water level data were obtained from the MLIT for the gauging station closest to the study segment at the downstream end (Kariki gauging station of the MLIT, ca.100 m downstream of the study segment).

### 2.5 Analyses

To characterize the spatial variation in HZ water quality in relation to tributary-derived GW influences, principal component analyses (PCA) were used to summarize the water quality data measured in the snapshot surveys. Six parameters were used in each of the winter and summer datasets: water temperature, pH, EC, differences in HZ water temperature relative to surface water (delta temperature), differences in HZ DO relative to surface water (delta DO), and differences in HZ EC relative to surface water (delta EC). Delta values were also included to highlight HZ characteristics with minimized effects of potential surface water longitudinal changes as well as diurnal changes in the environment (especially in temperature). A tributary-derived GW index was developed based on the extracted PCA axis scores for both seasons. In short, the scores of the axes, which were interpreted as an index of tributary-derived GW by referring to the results of our previous study (Negishi et al., under review), were used to represent the GW index at each point. To match the spatial scale of water quality data with salmon redds data, water quality was averaged to represent the 150-m compartment as analytical units hereafter.

To characterize temporal variation in HZ water, water quality data of HZ in intensive surveys were compared between selected 250-m sections by developing generalized linear (mixed) models (GLMs or GLMMs) separately for two seasons (Gaussian error distribution). GLMs with these three parameters as response variables were developed for EC, pH, and DO. For temperature, GLMMs with temperature as the response variable and measurement increments (every hour) were developed as random factors. To examine the temporal dynamics of tributary-derived GW in HZ water, continuous delta temperature data were plotted against the 14-day average water level.

The spatiotemporal dynamics of fish redds site selection were examined by developing generalized additive models (GAMs). As an explanatory variable, the survey month was included as the main explanatory factor, with the distance of respective compartments from the upstream and its interaction with survey months as a smooth term, and redds count as the response variable (negative binomial error distribution). The relative importance of environmental factors (GW index, channel slope, and distance to ground sluices) on redds site selection at the 150-m-conpartment scale was tested by developing GLMs separately for different seasons, with redds count as the response variable and three factors as explanatory variables (negative binomial error distribution). The variance inflation factor for the three parameters was <1.65, showing acceptably low levels of multicollinearity, and thus, all were included in the model after standardization.

The population-level consequence the presence of tributary-derived GW in HZ in the sea-migration salmon individuals in the following generation was examined by simulating redds HZ water temperature, and the time for egg development and the timing of fry emergence. First, the delta temperature was modelled in relation to the surface river water temperature at different GW index levels. Using the logger data obtained in the intensive monitoring survey on two seasons, a GLM was developed with delta temperature as a response variable, surface water temperature and the level of GW effects, and their interactions as explanatory variables (Gaussian error distribution). In this step, the level of GW effects (GW index) was estimated based on EC and linear regression models between EC and PC1 scores for each occasion developed in above-mentioned PCA analyses. Second, the modelled delta temperature was applied to continuous year-round surface water temperature data (average of 2001-2010 in MLIT) as well as GW index specific to each 150-m compartment so that HZ water temperature in each compartment was estimated (scenario with tributary-derived GW present in HZ). Another HZ water temperature scenario was modelled by applying the lowest GW index obtained to all 150-m compartments. Days until emergence since the survey days and fry emergence timing (Julian days) were modelled for each recorded redds (n=9699 redds for the period 2001-2015) in the respective scenarios based on their spawning location (compartment). Based on cumulative emergence (counted as redds number) over time, the Shannon-Weiner diversity index was obtained by considering redds spawned in different months as groups and comparing the temporal trends between the two scenarios. In these simulations, 980 degree-days were used as the timing of fry emergence.

All analyses, otherwise mentioned, were conducted using R (R Core Team, 2022) and relevant packages such as “*ggplot2*,” “*mgcv,*” and “*glmmTMB*.”

## 3. Results

The two PCA axes explained >70% of the variance in both the seasons (Table 1). Water quality parameters changed more longitudinally in the HZ than in the surface river water (Supplementary material 2). The first PC scores were similarly correlated with parameters in the raw and delta values of EC, pH, and DO. HZ water low in EC, DO, and pH was characterized by PC1. In PC1, the temperature correlation was inconsistent between seasons, with colder HZ water correlated with PC1 in summer, whereas its association was less pronounced and opposite in winter. In contrast, PC2 was inconsistently associated with the environmental factors during the two seasons. PC1 scores in the two seasons were indicative of similar HZ water quality and were spatially correlated with each other (Pearson’s correlation, *r*=0.69, *p*<0.001), indicating that HZ water in relatively downstream sections was characterized by unique water chemistry associated with GW (Fig. 2 a & b). Thus, the PC1 score in winter, which was available for all compartments in the study segment, was used as a tributary-derived GW index in subsequent analyses. GS distance tends to be shorter with more variable channel gradients in upper part of the segment (Fig. 2 c).

**Figure 2.**
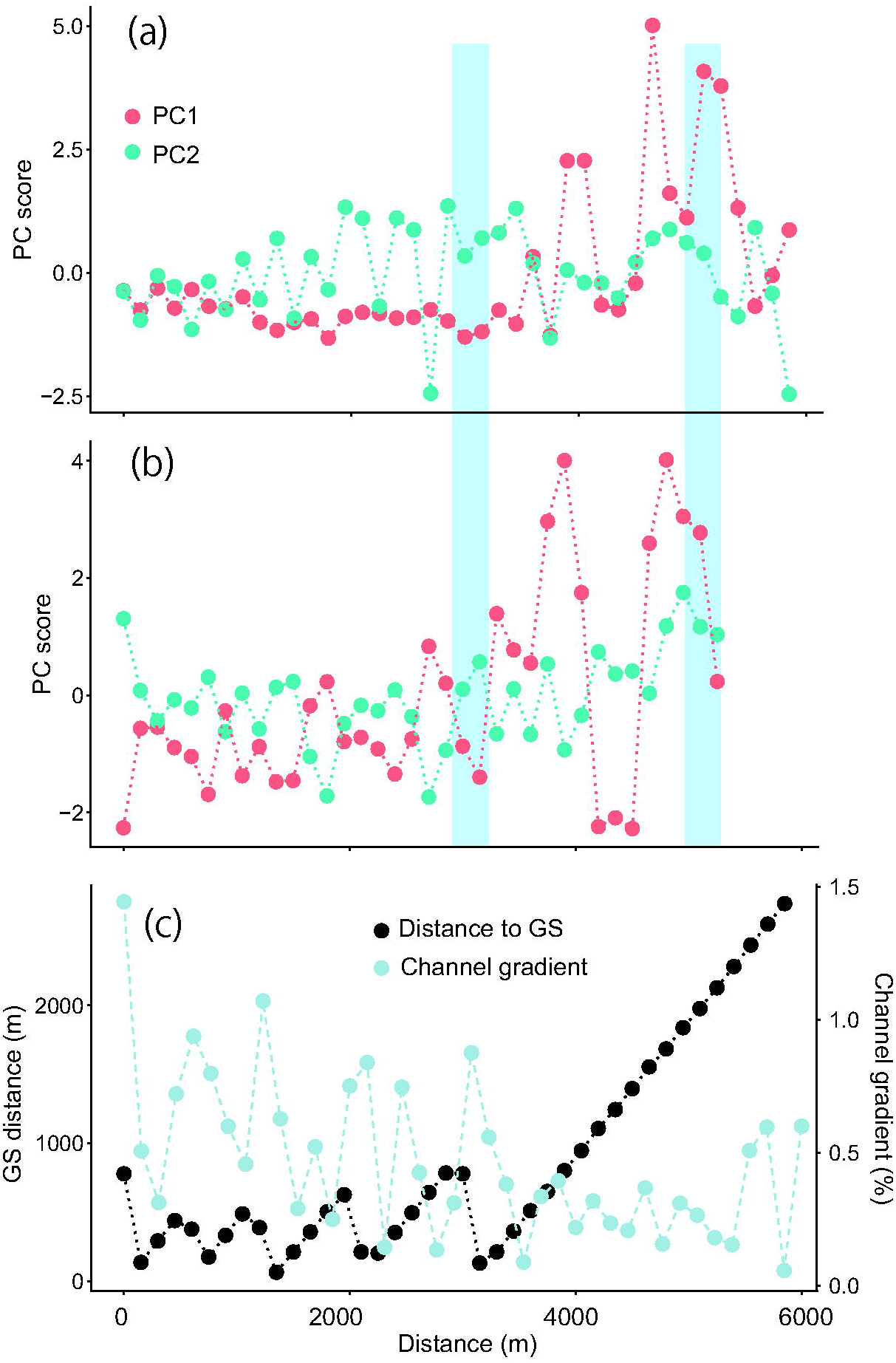
Spatial distribution of principal component scores in winter (a) and summer (b), and channel gradient and distances to immediate groundsills (c) in relation to the section’s distances from the upstream end of study segment. Blue bands shown in (a) and (b) denoted intensively monitored sections: upstream one for low tributary-derived groundwater effects (low GW section) and downstream one for high tributary-derived groundwater effects (high GW section).

**Table 1.** Results of principal component analysis to characterize hyporheic zone water (HZ water). Environmental factors are shown in relation to PC scores, with their eigenvalues and variance explained (%). The two seasons were analyzed separately. Abbreviations: environmental factors of HW: electrical conductivity (EC); dissolved oxygen (DO); temperature (temp); temperature differences of HZ water relative to surface water (delta temp); EC differences of HZ water relative to surface water (delta EC); pH differences of HZ water relative to surface water (delta pH); DO differences of HZ water relative to surface water (delta DO). Pearson’s correlation coefficients with the respective PC scores are shown with their statistical significance: *<0.05, **<0.01, and ***<0.001.

| Principal components | Winter |  | Summer |  |
| --- | --- | --- | --- | --- |
|  | PC1 | PC2 | PC1 | PC2 |
| Eigenvalues | 4.19 | 1.59 | 5.11 | 1.23 |
| Variance explained (%) | 52.85 | 20.06 | 64.56 | 15.54 |
| Environmental factor |  |  |  |  |
| EC | 0.88*** | 0.19 | 0.82*** | 0.48*** |
| pH | -0.79*** | 0.04 | -0.86*** | 0.26** |
| DO | -0.79*** | 0.46*** | -0.85*** | 0.31*** |
| temp | 0.04 | 0.67*** | -0.75*** | -0.15 |
| delta temp | 0.29*** | 0.80*** | -0.87*** | -0.23* |
| delta EC | 0.94*** | 0.17 | 0.81*** | 0.52*** |
| delta pH | -0.79*** | -0.21* | -0.57*** | 0.62*** |
| delta DO | -0.78*** | 0.45*** | -0.84*** | 0.34*** |

More intensive monitoring of HZ water in two 250-m sections with high PC1 scores (high GW section) and low PC1 scores (low GW section) were consistent with snapshot measurements. The water quality was more different between surface and HZ water in the high GW section (Supplementary material 3), showing a similar variation in relation to the GW level. The temperature difference was seasonally opposite in its direction, with HZ water being warmer in winter and colder in summer relative to surface water (Fig. 3 a & b), and with high EC and low DO in HZ water relative to surface water regardless of the season (Fig. 3 e‒h). The differences between section types were statistically significant (Table 2a). The water temperature difference between the HZ and surface water decreased in response to water level increases, at least in the summer (Fig. 3 c & d).

**Figure 3.**
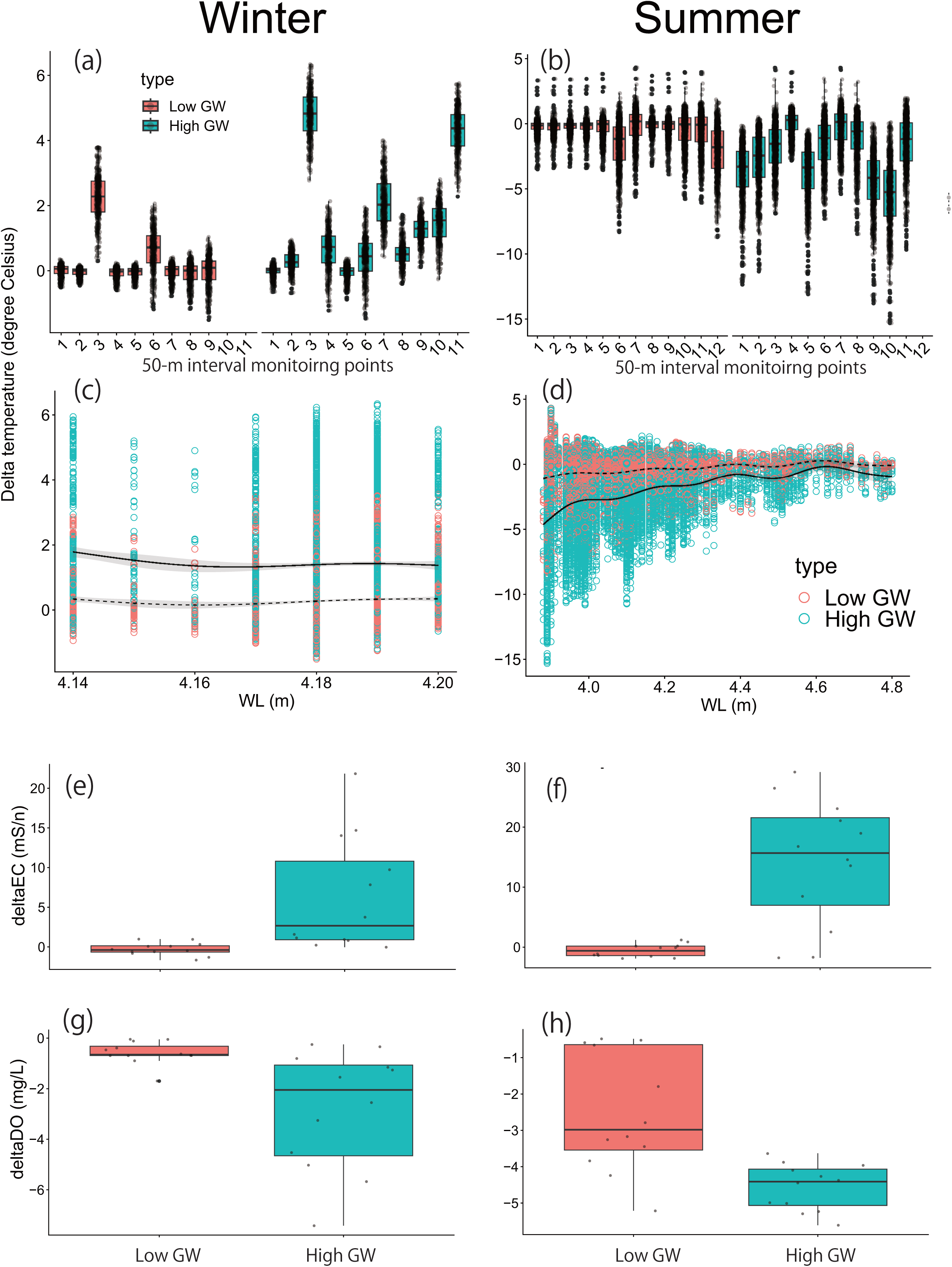
Box plots showing the differences (delta) in water temperature, EC, and DO of HZ water relative to surface water in winter (a, d, g) and in summer (b, f, h). Delta temperature changes in relation to water levels were also shown in winter (c) and summer (d). Solid (high GW section), broken lines (low GW section), and gray clouds in (c) and (d) denote statistically significant regression lines developed by generalized additive models (water level as a main smoothing factor with Gaussian error distribution) and its 95% confidence intervals for each HZ water type.

**Table 2.** Results of generalized (mixed) linear models (GLMs or GLMMs) testing the effects of section types (high groundwater and low groundwater sections) on differences (delta) in water temperature, EC, and DO of HZ water relative to surface water in different seasons (winter in February and summer in August) (a), and testing the effects of groundwater (GW index), channel slope, and distance to groundwater (GS distance) on fish redds counts in different seasons. In (b), the parameters are standardized prior to the analyses. The standard errors are shown in parentheses. Abbreviations: water temperature (temp), electrical conductivity (EC), dissolved oxygen (DO), September (Sep), October (Oct), November (Nov), December (Dec), and January (Jan).

| (a) |  |  |  |
| --- | --- | --- | --- |
| Parameters | Estimates | Z | p |
| Delta temp (winter) | 1.13 (0.03) | 40.01 | <0.001 |
| Delta temp (summer) | -1.87 (0.02) | -89.27 | <0.001 |
| Delta EC (winter) | 6.66 (2.01) | 3.31 | <0.001 |
| Delta EC (summer) | 14.83 (2.91) | 5.10 | <0.001 |
| Delta DO (winter) | -2.23 (0.66) | -3.37 | <0.001 |
| Delta DO (summer) | -2.07 (0.49) | -4.24 | <0.001 |
| (b) |  |  |  |
| GW index (Sep) | -0.02 (0.12) | -0.19 | 0.51 |
| Slope (Sep) | -0.41 (0.11) | -3.60 | <0.001 |
| GS distance (Sep) | -0.58 (0.14) | -4.08 | <0.001 |
| GW index (Oct) | 0.11 (0.06) | 1.75 | 0.08 |
| Slope (Oct) | -0.45 (0.07) | -6.73 | <0.001 |
| GS distance (Oct) | -0.44 (0.08) | -5.73 | <0.001 |
| GW index (Nov) | 0.36 (0.08) | 4.45 | <0.001 |
| Slope (Nov) | -0.81 (0.08) | -9.58 | <0.001 |
| GS distance (Nov) | -0.39 (0.09) | -4.33 | <0.001 |
| GW index (Dec) | 0.40 (0.10) | 4.01 | <0.001 |
| Slope (Dec) | -1.10 (0.13) | -8.69 | <0.001 |
| GS distance (Dec) | -0.14 (0.11) | -1.30 | 0.19 |
| GW index (Jan) | 0.18 (0.14) | 1.25 | 0.21 |
| Slope (Jan) | -1.23 (0.20) | -6.03 | <0.001 |
| GS distance (Jan) | -0.16 (0.17) | -0.94 | 0.35 |

The salmon redds counts were distributed along the river channel in relation to month and distance (Table 3). October redds counts were highest, with the lowest in January, and the seasonal longitudinal distribution varied (Fig. 4). There was a relative peak of redds selection in the upper part of the segment (distance of approximately 2,000 m) in the early months, and this peak became less pronounced in December and January. In contrast, the lower section was less pronounced in September (distance of 4,000‒5,000 m) and became abundant in redds count, starting from October to December. The relative importance of different environmental factors in explaining redds selection differed among months (Table 2b; Supplementary material 4). The effects of the channel slope were consistently negatively significant, whereas the GW index had significant effects in November and December. The distance to the groundsills was important in the early months but became insignificant in December and January.

**Figure 4.**
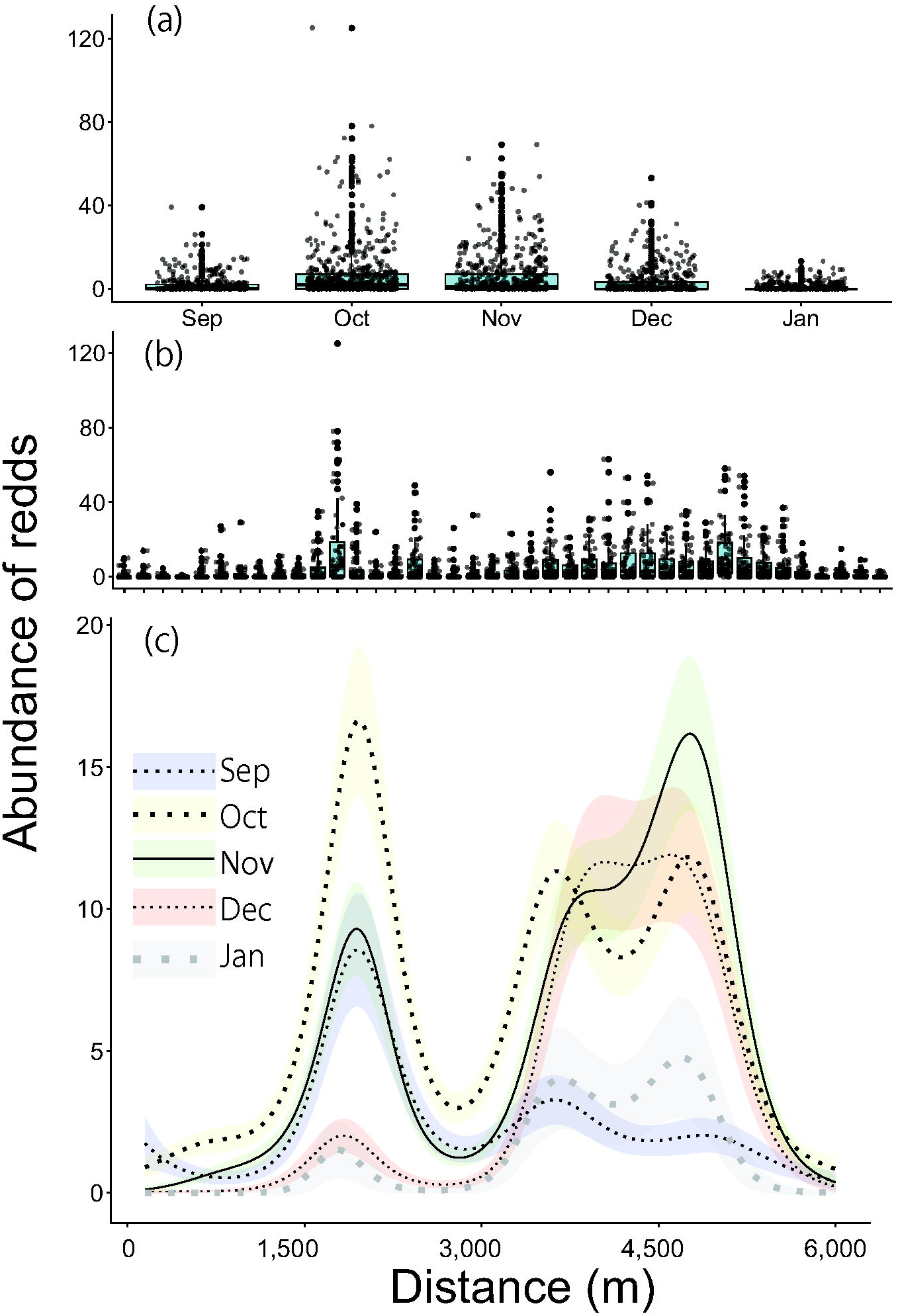
Box plots showing salmon redds counts in different month (a), 250-m survey compartment (b), and model regression based on generalized additive models (survey month as a categorical term, the interaction between distance to the upstream end and survey month as a main smoothing factor) (c). In (b), each group is arranged according to the compartment’s distance from the upstream end of the study segment; in (c), the distance is expressed as a continuous distance value from upstream to downstream. Abbreviations: September (Sep), October (Oct), November (Nov), December (Dec), and January (Jan). Model lines were described for different survey months, with clouds around the lines denoting 95% confidence intervals for each relationship.

**Table 3.** Results of the generalized additive model (GAM) for variations in fish redds count in each 150-m reach section for each month responding to interactions between distance and month. The estimates for the smooth terms are the estimated degrees of freedom. Standard errors for categorical terms are shown in brackets, and categorical terms were developed as groups against data in December. Abbreviations: September (Sep), October (Oct), November (Nov), December (Dec), January (Jan), and distance from the upstream end (Dis). Smooth terms are interactions between the month and distance from the upstream end. †Statistics for categorical terms and smooth terms are Z statistics and chi-square statistics.

| Terms | Estimates | Statistics<sup>†</sup> | *p* | Explained variance | R<sup>2</sup> |
| --- | --- | --- | --- | --- | --- |
| Categorical terms |  |  |  |  |  |
| Intercept | 0.08 (0.11) | 0.79 | 0.43 |  |  |
| Dec-Jan | -2.52 (1.15) | -2.19 | <0.05 |  |  |
| Dec-Nov | 1.00 (0.12) | 8.57 | <0.001 |  |  |
| Dec-Oct | 1.46 (9.11) | 12.97 | <0.001 |  |  |
| Dec-Sep | 0.51 (0.12) | 4.15 | <0.001 |  |  |
| Smooth terms |  |  |  |  |  |
| Dis:Dec | 8.30 | 585.09 | <0.001 |  |  |
| Dis:Jan | 6.95 | 66.34 | <0.001 |  |  |
| Dis:Nov | 8.71 | 752.91 | <0.001 |  |  |
| Dis:Oct | 8.79 | 606.36 | <0.001 |  |  |
| Dis:Sep | 8.40 | 192.76 | <0.001 | 40.30 | 0.21 |

<sup>†</sup>Statistics for categorical terms and smooth terms are Z statistics and Chi-squared.

The Delta temperature was statistically significantly modelled by surface water temperature, GW index, and their interactions (Supplementary material 5). HZ water is generally warmer and colder than the river water in winter and summer, respectively. This deviation from the surface water increased with increasing GW effect. Based on the model, HZ water in each compartment within the study segment showed spatially heterogeneous water temperature trajectories over time, whereas low GW-scenario simulations were spatially single and only slightly deviated from the surface river water temperature (Fig. 5). Both emergence days and timing of emergence were more heterogeneous and extended to shorter days (for emergence days) and earlier Julian days (for emergence timing) when GW effects were present (Fig. 6 a–d). Over the entire emergence period, fry emergence groups regarding their spawned timing remained more diverse when GW effects were present (Fig. 6 e & f).

**Figure 5.**
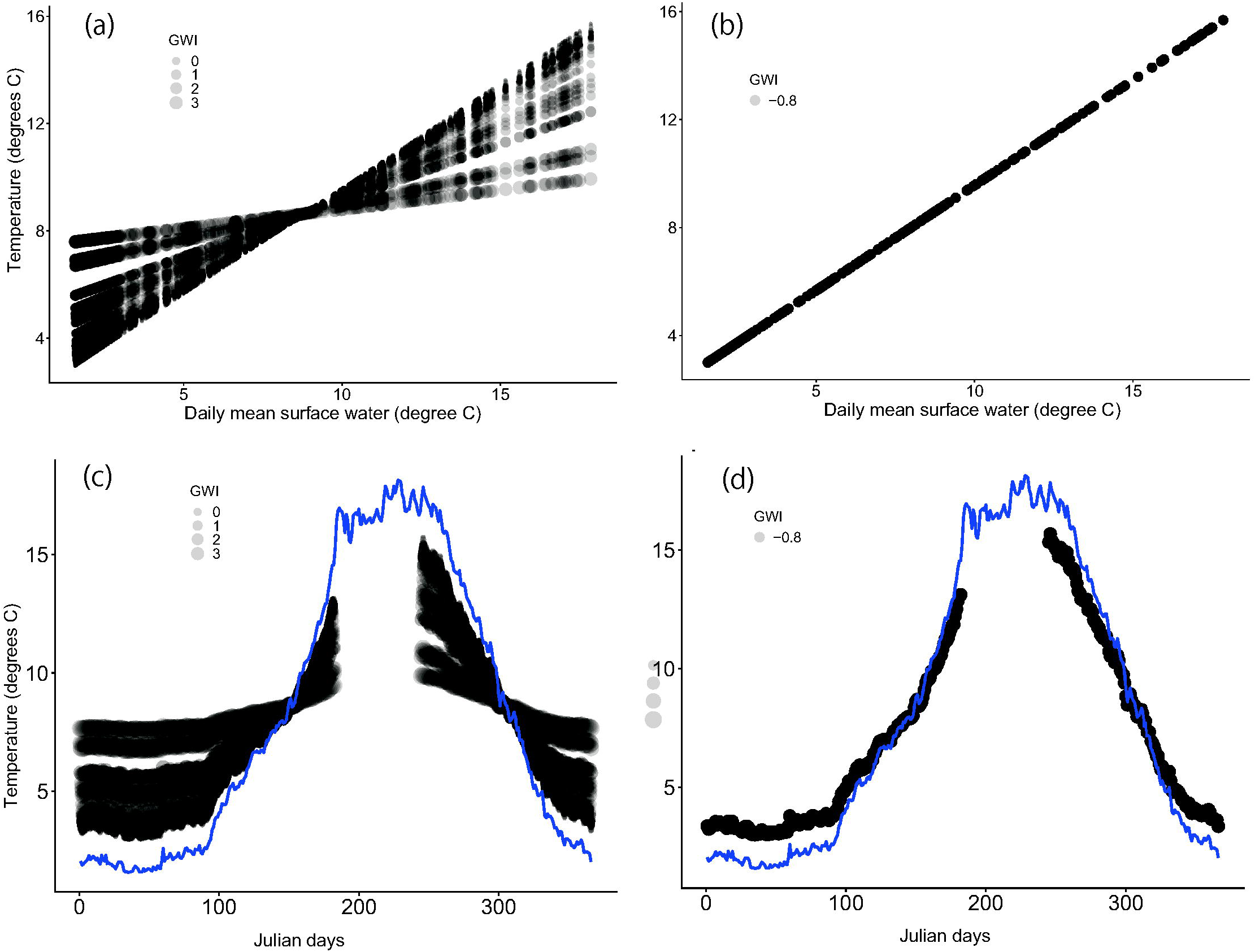
Predicted HZ water temperature in relation to surface water temperature study sections with (a) and without groundwater effects (b) for the respective survey compartments. Simulated HZ water temperature in relation to observed surface water temperature with (c) and without groundwater effects (d) for respective compartments. The groundwater index (GWI) is shown as the size of the dots. The blue lines in the lower panels denote the surface water temperatures observed in 2001‒2010. The modeled temperature was absent for the middle summer because no simulation was conducted for these time windows to save computational resources.

**Figure 6.**
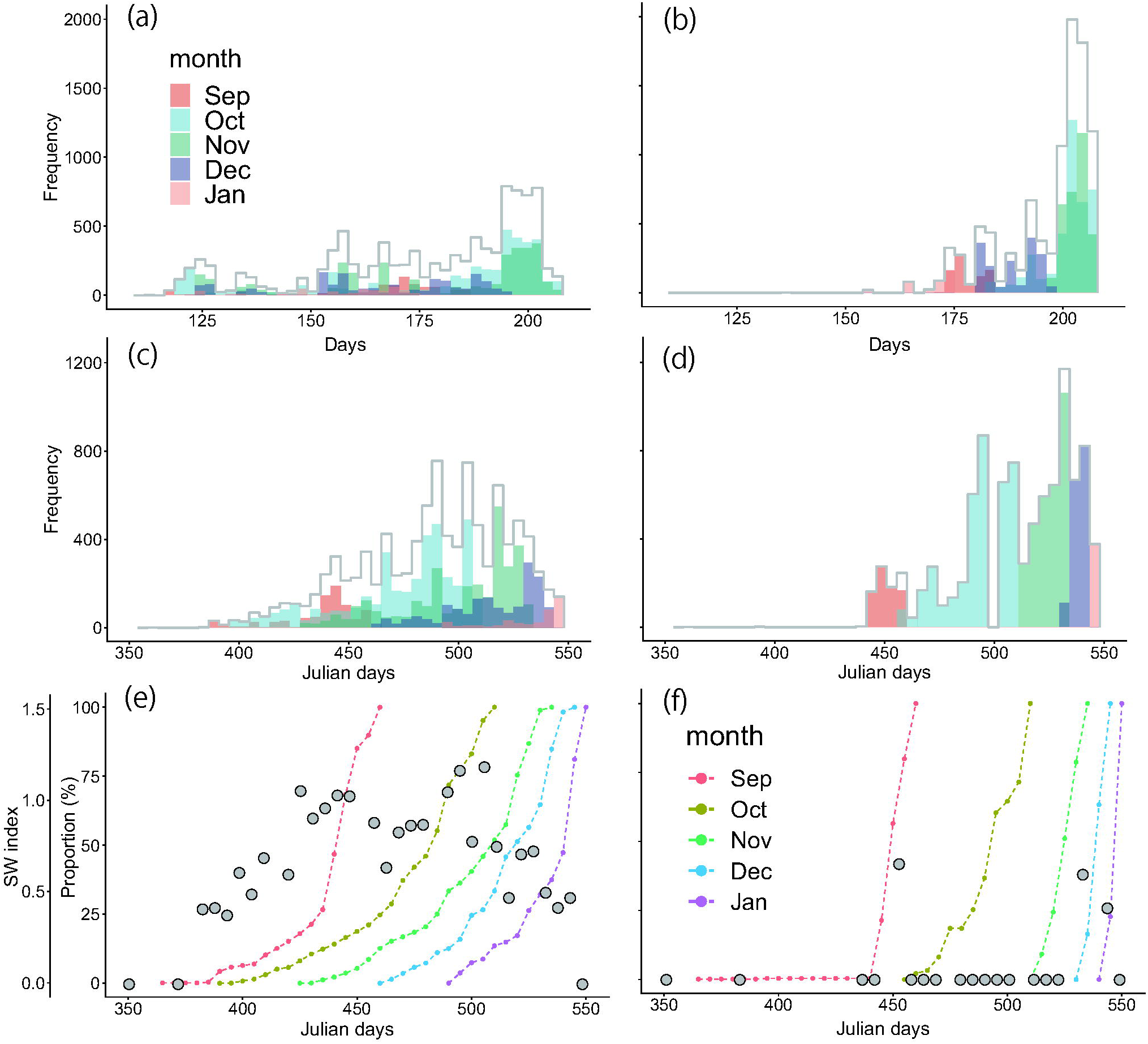
Simulated days required from spawning until fry emergence according to their redds formation with (a) and without (b) groundwater (GW) effects. Julian days for the timing of fry emergence according to their redds formation with (c) and without (d) GW effects. Counts are shown for monthly groups to highlight temporal variations in the responses of spawning individuals. Shannon-Wiener index (SW index) based on groups according to their redds formation and cumulative proportion of fry from groups at respective Julian days with (e) and without (f) ground water effects. The gray outlines in the upper four panels denote the total frequency (individuals), including all monthly groups. Note that the Julian days in the lower four panels were continuous from the previous year; the 366th day was 1st January.

## 4. Discussion

The predictions that salmon redds site selection is affected by the presence of groundwater and that its importance becomes more pronounced in winter periods were both quantitatively supported by the findings. Modelling exercises also revealed that this seasonally variable redds formation in relation to groundwater would manifest as population-level consequences in their offspring’s development and timing of descent to the sea in the following year. These findings extend those of previous studies that associated groundwater-affected habitats for redds selection of salmon at different spatial scales (Mouw et al., 2014; Aruga et al., 2023) and highlighted their importance for anadromous salmon, even within a segment <10 km. The present study, based on a coupled understanding of hydrochemical processes and biological responses, demonstrated that the importance of thermally diverse redds sites, as hot spots for spawning salmon in harsh cold winters, can be recognized in the early stages of salmon hatching in freshwater environments from the perspective of a well-adapted species strategy to uncertain environments.

Groundwater-affected hyporheic zones with unique characteristics relative to the surface water were identified in the lower part of the study segment, as anticipated. These areas were consistently characterized by HZ water having higher EC and relatively lower DO, and distinct temperatures (colder in summer and warmer in winter) relative to surface water. Water temperature did not appear to be a strong parameter for winter GW-affected HZ water, but this was not surprising because surface water warmed gradually during the field work, which coincided with the surveys in the GW-affected section.

As a result, the difference in temperature diminished because it was when the water temperature in both zones became similar to each other at approximately 10 degree C, resulting in little temperature difference between the two zones. However, continuous monitoring of the sections in the colder months (February) provided results consistent with the notion that groundwater also provides warm water in winter during colder periods (Power et al., 1999). It is known that groundwater upwelling from the riverbed can be chemically and thermally different depending on their sources, associated flow pathways, and residence time, with deeply and remotely originated groundwater that often has more chemically unique properties (Alexander & Caissie, 2003).In a parallel study, Negishi et al. (under review) revealed that this groundwater originates from a tributary catchment and continues to subsidize the food webs of the river and riparian communities in the study segment. Overall, the present study added ecological functions to this subsurface connectivity in terms of controlling salmon spawning habitat quality.

The GW-affected HZ water was warmer in winter, and salmon spawning beds tended to be formed in sections dominated by warmer areas. This trend of redds site selection preference of GW-affected water areas in colder seasons is consistent with the findings of Aruga et al. (2023). Several important implications for GW properties and salmon usage as a thermally distinct spawning zone can be provided based on the new knowledge synthesized in these two related studies. Aruga et al. (2023) demonstrated that redds were distributed seasonally according to geomorphic units, earlier in upwelling areas of riffles, and later in secondary channels. We found that GW presence was not restricted to geomorphic units per se, but was longitudinally distributed, including the river bed in the main channel, because our methods represented all different types of geomorphic units along the thalwag. Thus, this collectively indicates that for the GW identified in the present study, geomorphic units are not the cause or direct areas affected by GW, but such geomorphic units occur at least partially in the sections where GW upwelling dominates. The important information gap of Aruga et al. (2023) in the discussion of groundwater nature was the lack of simultaneous measurements of chemical properties, such as EC, which was clearly found to be the key parameter in appreciating the origin and identity of GW. As a result, warmer redds associated more frequently with secondary channels in later seasons could have included those affected by the GW (the same as ours) and GW sourced from nearby areas, such as riffle heads only distinct in water temperature (Brunke & Gonser, 1997; Baxter & Hauer, 2000). The latter was not targeted in the present study, and such areas might have caused some spotty occurrence of high PC values in upstream areas. Thus, it is possible that the present study ignored other types of GW-affected HZ, indicating that several types of GW-affected spawning redds may exist. Overall, the two studies agree on the presence of thermally diverse habitats within a relatively small <10 km segment.

The importance of GW-affected thermal heterogeneity at different spatial scales and in different geographical areas has been well discussed in the context of fry emergence timing and salmon descent into the sea. However, such variations have been empirically examined or modelled in a simulative framework for surface water, largely neglecting the HZ temperature and its spatial variations in the difference from surface river water. For example, Crozier et al. (2021) modelled population dynamics, including fry emergence timing at different temperatures, using surface-water modeling. Other studies have also reported variations in salmon runs and sea descends of fry in relation to variations in surface water among sites (Webb & McLay, 1996; Lisi et al., 2013). Aruga et al. (2023) preceded the present study to report on the possible relationships between spatial thermal heterogeneity and potential variations in egg development and fry emergence in the context of GW-affected HZ water, but were unable to estimate how such diversity may lead to population-level consequences. The largest source of hindrances to such efforts was the lack of HZ water temperature data encompassing incubation periods and the range of GW influences. The present study is the first to take this step forward and successfully show how thermally diverse HZ water may result in the diversification of timing of early salmon life stage. GW-affected HZ water allows the formation of heterogeneous thermal templates and thus spatially dynamic redds formations, which in turn lead to more synchronized descending timing of fish from eggs laid at different times of the previous year. The selection of such a strategy was consistent with the expected traits of adaptation to natural environments. Different strains (spawners at different times; early– and late-run individuals; sensu Aruga et al., 2023) can have a better chance of coping with uncertain environmental conditions in their descends; if descending in a sequential manner according to spawning timing, the risk of high mortality concentrated on specific groups associated with sudden events will be high (the second scenario).

There are several caveats that must be taken into account when making interpretations of results for salmon habitat management. First, the GW quality in this study was probably not optimal for salmon egg development because it is characterized by low DO. We suspect that together with exceedingly high EC values, the effects of human activities that pollute its sources or water along its pathway to the river exist in contemporary urbanized settings (Negishi et al., under review). Thus, potential negative effects such as increased mortality rates of fry from GW-affected redds need to be incorporated to obtain better estimates of GW effects on salmon population dynamics and highlight the need for further experimental testing on physiological responses to this groundwater. Second, our modelling approach did not use data of temperature and water quality collected in real redds as previously done in Aruga (2023), and thus, the model calibration with the real data also uses fry emergence timing and temporal changes of fry abundance is preferable. We observed in parallel studies that spawning redds formed in late seasons were characterized by the GW index and temperature falling within the range of the GW index and thermal regime reported in this study. Furthermore, we used 980 degree-days as the timing of fry emergence, but this criterion may be different in the field environment and could be an overestimate for populations in the study area (Ohmoto, 2018). An improved understanding of the life-cycle timings specific to the Toyohira River is needed for more accurate models. Third, our estimates were based on redds counts, ignoring individual variety in mortality, redd-specific mortality rates, and redd-specific egg abundance variations. Despite of some limitations, we believe that the general trends found and the interpretations of the data in this study were acceptable.

In conclusion, the present study identified hotspots for salmon spawning redds in winter with a disproportionately high level of site selection because of its warmer temperature compared to surface river water in winter. Thermally diverse spawning habitats allow the diversification of spawner strains in synchronized descents to the sea. Signs of GW pollution were suggested, and thus the conservation of the groundwater pathway and its sources, followed by improvements in quality, can be beneficial to the Chum salmon populations in the Toyohira River. The presence of GW zones and their biological importance have been well conceptualized and known, but empirical data to clarify such associations are still scarce. The presence of such thermally diverse environments could enhance the resilience of fish populations in the face of environmental changes because of the diversification of development and behavior at the individual and population levels. The effects of climate change on flow regimes in winter as well as groundwater are of concern (Jyrkama & Sykes, 2007; Isaak et al., 2012), and this is alarming because the relationship between water level and thermal characteristics of GW-affected HZ water was also found. Climate change effects may directly or indirectly mask the thermal heterogeneity associated with GW sensed by animals in a complex manner. Although summer maxima tend to receive attention for river conservation in the context of a changing climate, hot spots in winter also need to be identified and analyzed in species sensitivity studies.

## Supporting information

Supplementary materials all

## Acknowledgements

This study was partly supported by the research fund for the Tokachi and Ishikari rivers provided by MLIT (18056588) and JSPS KAKENHI (18H03408 and 18H03407).

## Ethical approval

No ethical violation was occurred in this research.

## Data availability statement

All the data used in the paper will be deposited in Dryad.

## Author contributions

JN Negishi: Conceptualization (equal), data collection (equal), data curation (equal), formal analysis, funding acquisition (equal), and leading writing. N Morisaki and YY Song: Conceptualization (equal), data collection (equal), data curation (equal), and preliminary analyses (equal). N Aruga: Conceptualization (equal), data collection (equal), and data curation (equal): H Urabe and F Nakamura: Conceptualization (equal), data curation (equal), and funding acquisition (equal). All the authors contributed critically to the drafts and approved the final manuscript for publication.

