## Supplementary materials all for "Hot spots in a cold river synchronize temporarily separated salmon runs in their offspring’s development"

* Corresponding author

**Supplemental material 1**

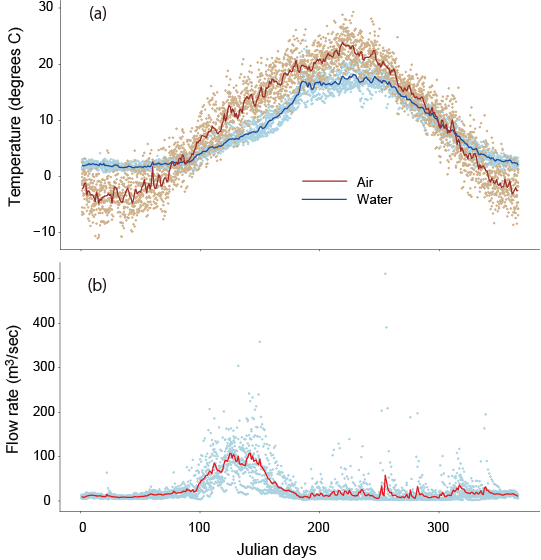

Air and water temperature (a) and flow rate (b) of the Toyohira River measured in the period 2001-2010 at Horohirabashi Gauging station of the MLIT, which is approximately 1 km upstream of the study segment. Daily average values are plotted with their means in solid lines.

**Supplemental material 2**

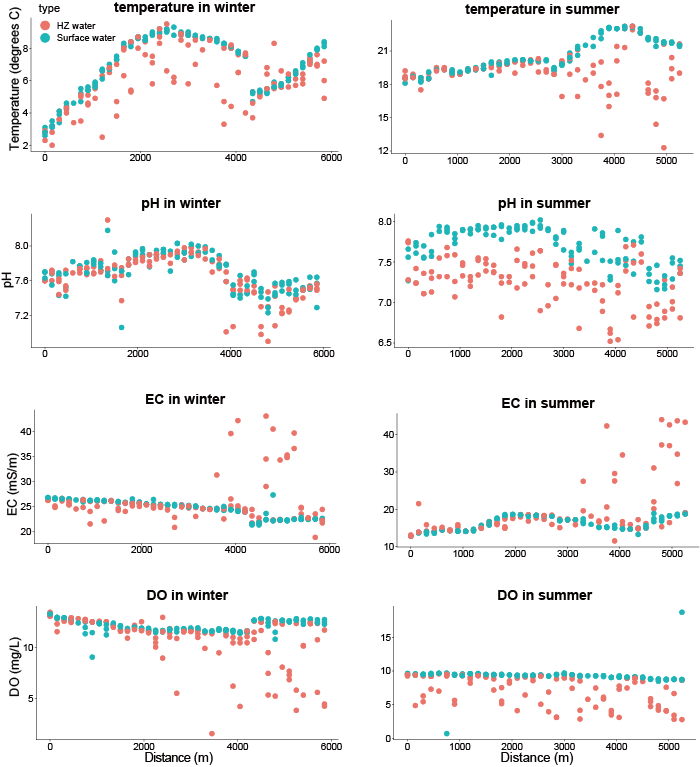

Longitudinal trends of water quality parameters in the hyporheic zone (HZ) and surface water in winter and summer. The horizontal axis represents the channel distance measured from the upstream end of the study segment.

**Supplemental material 3**

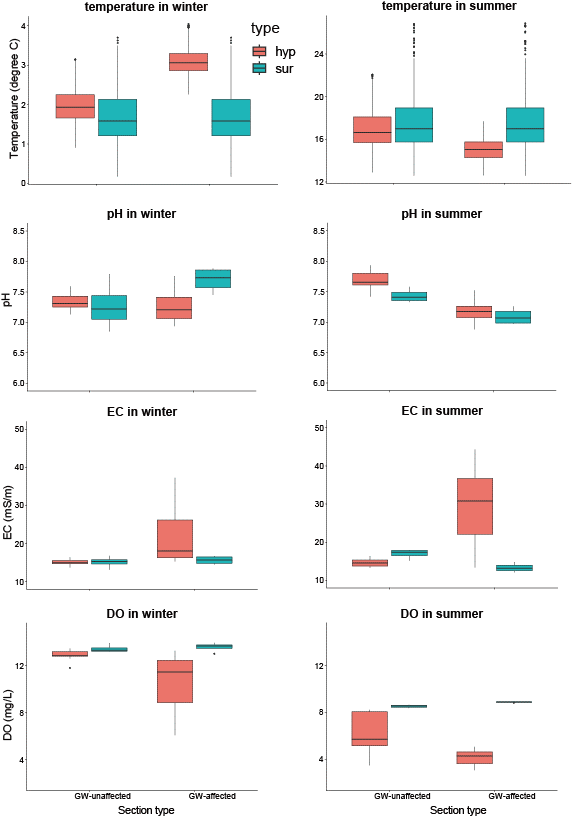

Trends of water quality parameters in hyporheic zone (HZ) and surface water in winter and summer in GW-affected section and GW-unaffected section.

**Supplemental material 4**

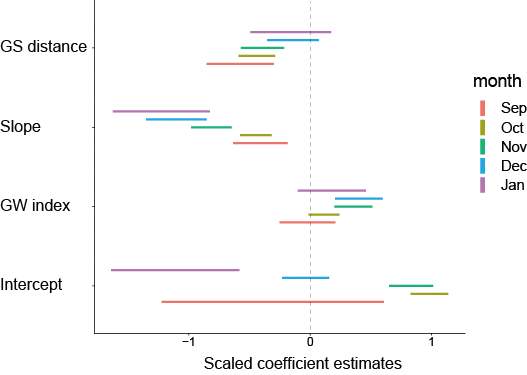

Coefficients in generalized linear models testing the effects of groundwater (GW index), channel slope, distance to ground sills (GS distance) on fish redds count in different seasons.

**Supplemental material 5**

| Parameters | Estimates | Z | *p* |
| --- | --- | --- | --- |
| Intercept | 2.88 (0.35) | 8.20 | <0.001 |
| Temp | -0.35 (0.01) | -24.10 | <0.001 |
| GWI | 1.39 (0.36) | 3.89 | <0.001 |
| Temp × GWI | -0.16 (0.01) | -10.51 | <0.001 |

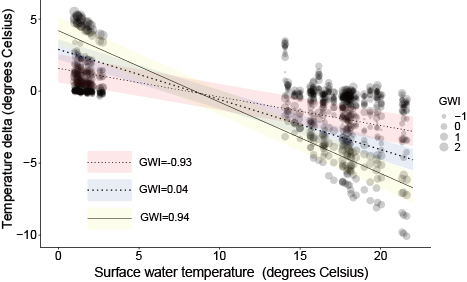

The table shows the results of a generalized mixed linear model testing effects of surface water temperature (Temp) and groundwater index (GWI) and their interactions on water temperature difference of hyporheic water relative to surface water temperature. Standard errors were shown in brackets. The lower panel shows data with modelled relationships; modelled relations were shown for three levels of GWI as examples.
